# A tool to dissect heterotypic determinants of homotypic protein phase behavior

**DOI:** 10.1101/2025.01.01.631016

**Authors:** Hannah Kimbrough, Jacob Jensen, Caleb Weber, Tayla Miller, Lucinda E. Maddera, Vignesh Babu, William B. Redwine, Randal Halfmann

## Abstract

Proteins commonly self-assemble to create liquid or solid condensates with diverse biological activities. The mechanisms of assembly are determined by each protein’s sequence and cellular context. We previously developed distributed amphifluoric FRET (DAmFRET) to analyze sequence determinants of self-assembly in cells. Here, we extend DAmFRET by creating a nanobody (mEosNb) against the fluorescent protein mEos3 to physically tether and thereby recruit candidate modifier proteins to mEos3-fused query proteins. This accessorization allows us to rapidly screen for effects on the phase behavior of query proteins by modulating the expression level and valency of mEosNb-fused modifiers. We show that our system recapitulates known effects of multivalency on liquid-liquid phase separation and can discriminate between nucleation mechanisms of amyloid and amyloid-like assemblies. Our approach adds a new experimental dimension for interrogating the mechanisms of intracellular phase transitions.

**Lay summary:** Protein self-assemblies are essential for cellular function, but can also contribute to disease. We develop a new tool to study how their formation is influenced by other cellular factors. This tool allows us to control the location and number of interactions between a protein of interest and other proteins that may influence it. Our results provide new insight into mechanisms of self-assembly and will aid research toward treating diseases associated with aberrant assembly.

## Introduction

Proteins commonly self-assemble into liquid or solid phases with important biological consequences. Disordered liquid-liquid phase separation (LLPS) dynamically regulates and localizes protein activities (Alberti and Hyman, 2021; Lyon *et al*., 2021), while ordered less-dynamic phases including amyloid fibrils function as scaffolds, molecular memories, and switches that determine cell fate (Shi *et al*., 2020; Rodríguez Gama *et al*., 2021; Sawaya *et al*., 2021; Buchanan *et al*., 2023). Pathological amyloids also form in the course of Alzheimer’s and other progressive degenerative diseases as a consequence of age- or stress-driven solidification of initially dynamic protein assemblies. Hence, mechanisms of assembly govern the function or dysfunction of phase separation. These mechanisms are in turn determined by the interplay of each protein’s sequence and cellular context.

Mechanistic differences manifest most in the initial formation, or nucleation, of condensed phases. LLPS involves only a change in intermolecular density, while amyloid involves changes in both intermolecular density and intramolecular ordering (conformation). The latter renders the nucleation of amyloids less sensitive to concentration, and therefore probabilistic. Amyloid-forming proteins can consequently accumulate to highly supersaturating concentrations while remaining (temporarily) soluble, providing a thermodynamic drive that makes their formation effectively irreversible.

We previously developed an assay to characterize mechanisms of protein self-assembly in cells by evaluating their different concentration dependencies and kinetics. In this assay, known as distributed amphifluoric Förster resonance energy transfer (DAmFRET), proteins or mutants of interest are expressed in yeast or human cells as single genetic fusions to the photoconvertible fluorescent protein mEos3 (Khan *et al*., 2018). Following en masse limiting photoconversion, cells are analyzed by flow cytometry to quantify ratiometric FRET between the green and red forms of mEos3 (AmFRET) as a function of each protein’s intracellular concentration, taking advantage of heterogeneous expression levels between cells in the culture. The duration of protein expression (hours) in a DAmFRET experiment is long relative to timescales of LLPS, but short relative to timescales of amyloid nucleation in vivo. Consequently, LLPS manifests as a continuous acquisition of AmFRET beyond the protein’s saturating concentration (C_sat_), while amyloid formation tends instead to manifest as a discontinuous acquisition of AmFRET. Hence, DAmFRET facilitates comparative assessments of proteins to uncover sequence determinants of self-assembly (Posey *et al*., 2021; Kandola *et al*., 2023).

Myriad cellular factors also influence phase behavior in trans through diverse equilibrium and kinetic mechanisms, including mass action solubilization, complex coacervation, emulsification, and heterogeneous nucleation (Konstantoulea *et al*., 2021; Villegas *et al*., 2022). The effects of direct interactors strongly depend on their valence, or number of noncompeting quasi-equivalent binding sites for the phase separating protein. For LLPS, low-valence interactors tend to solubilize (raise C_sat_) while high-valence interactors tend to promote assembly (lower C_sat_) (Banani *et al*., 2016; Pak *et al*., 2016). Multivalent interactors can also accelerate the kinetics of ordered phase transitions by stabilizing oligomers at or beyond the size required to template subunit addition (Buell, 2017).

Mechanisms of trans-acting factors can be difficult to discriminate experimentally, however, and more tools are needed to facilitate such efforts. DAmFRET can measure the effects on self-assembly of proteins expressed in trans, but it presently provides limited information on the mechanisms of those effects. For example, the protein may facilitate assembly by binding the query protein, or instead by competing for a solubilizing factor such as a molecular chaperone. Conversely, a protein with relevant biochemical activity may fail to have an effect because it localizes to a different subcellular compartment.

Adapting DAmFRET to more effectively study trans-acting factors will require experimental control over the expression, valence, and physical proximity of those factors relative to the mEos3-fused query protein. Achieving such control would unlock a new experimental dimension for the hundreds of proteins that have already been cloned as fusions to mEos3 ((Khan *et al*., 2018; Posey *et al*., 2021; Gama *et al*., 2024); and unpublished).

The most direct way to physically link two proteins is by genetic fusion. This approach becomes cost prohibitive, however, for more than a small number of protein pairs. It also undesirably fixes their relative expression level, which is an important experimental variable for probing the mechanism of effect. For example, a stoichiometric effect of a modifier on phase separation would suggest that it (de)stabilizes the query protein assembly by directly binding it, whereas a substoichiometric effect would suggest that it instead acts enzymatically or as a nucleating template.

To facilitate the study of trans-acting modifiers of phase separation, we here accessorize DAmFRET by developing an mEos3-binding reagent to physically tether candidate factors to mEos3-fused query proteins at variable stoichiometry. Our system involves a custom-made nanobody against mEos3 that, when genetically fused to candidate factors and co-expressed with mEos3-fused query proteins, allows for the effects of interaction on self-assembly to be surveyed via flow cytometry for large numbers of query and modifying proteins across a wide range of relative expression levels.

## Results

### mEosNb discovery and validation

To develop an mEos3-binding reagent for intracellular expression, we employed a yeast surface display library of more than 1 x 10^8^ unique nanobody clones (McMahon *et al*., 2018). Derived from camelid antibodies, nanobodies (Nb) are much smaller than conventional antibodies (15 kD compared to 150 kD), do not provoke an immune response, and stably fold and function inside cells. From two independent screening campaigns against purified mEos3.1, we obtained a total of 6 unique clones, of which two exhibited good affinity based on on-yeast EC50 determination (not shown). The higher affinity clone, hereafter designated “mEosNb”, was identified in both screens. We used biolayer interferometry to accurately measure its affinity against purified mEos3.1, obtaining a dissociation constant (Kd) of 76.84 +/- 11.33 nM (**Figure 1A**).

**Figure 1.**
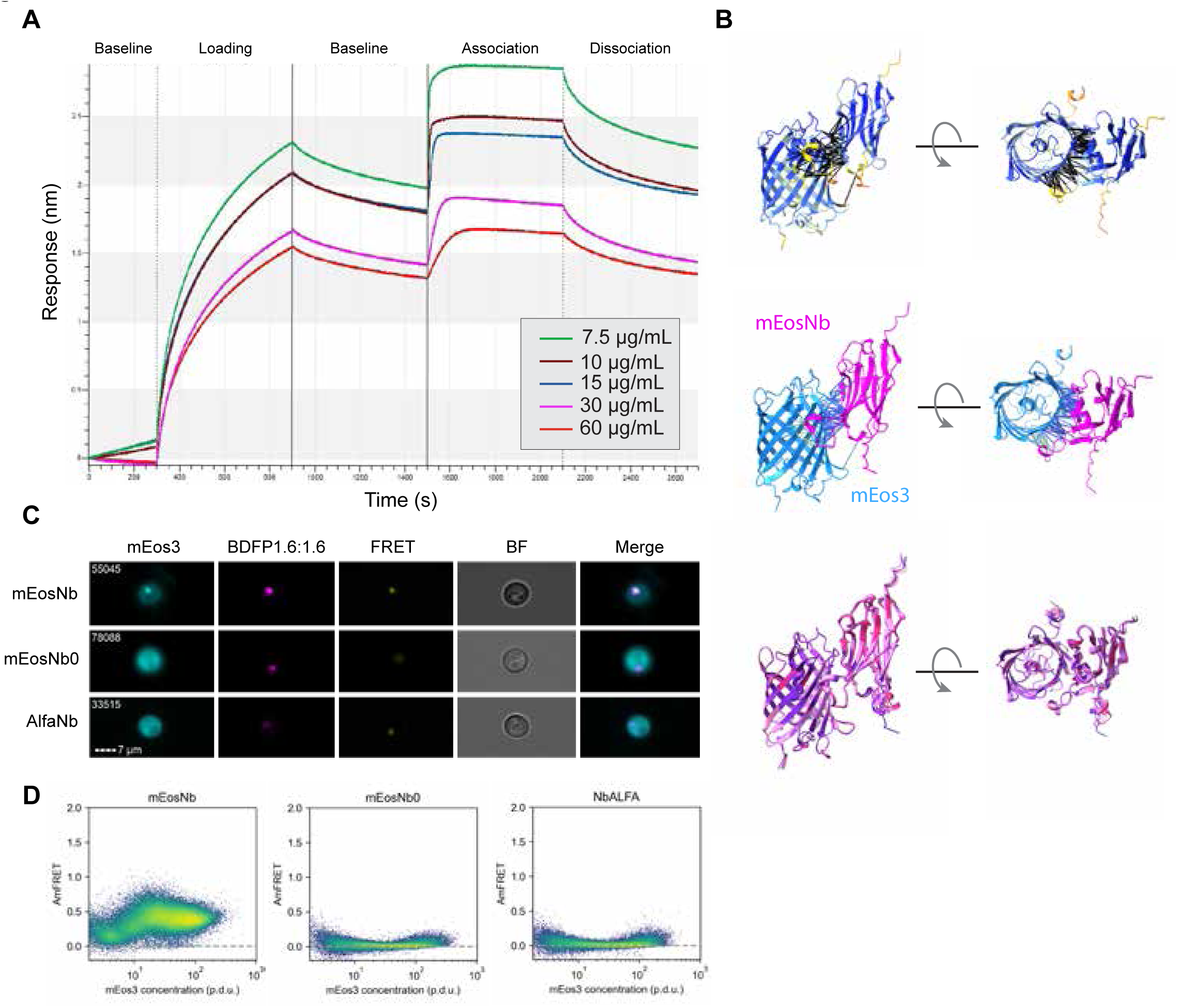
mEosNb discovery and validation. (a) Biolayer interferometry traces of Nb candidates binding to purified mEos3.1. (b) AlphaFold3 structural predictions of mEos-mEosNb complex colored by pLDDT (top) or by protein (middle), and of all five model predictions after structural alignment (bottom). Contact residues (middle) are predicted at 5Å and colored by PAE score. (c) Nanobody integrated yeast strains expressing (GA)20 amyloid-forming peptide assessed for subcellular colocalization using imaging flow cytometry and (d) self-assembly via DAmFRET.

We predicted the structure of the mEos3-mEosNb complex using AlphaFold 3 (Abramson *et al*., 2024). All five top scoring models were in excellent agreement, yielding an unambiguous structure (high pLDDT, low PAE, and Cα RMSD for all residues ranged from 0.386 to 2.627Å; **Figure 1B**) with an extensive interface involving 20 residues of mEosNb, consistent with its high affinity.

To test if mEosNb binds mEos3 in living cytoplasm, we created yeast strains expressing either mEosNb or the alternative clone (“mEosNb0”) identified in our screens, or as a negative control, a nanobody to ALFA-tag (Götzke *et al*., 2019). Each Nb was fused on its C-terminus to both µNS 471-721, which forms dense puncta in cells (Schmitz *et al*., 2009), and BDFP1.6:1.6, a far red fluorescent protein (Ding *et al*., 2018) that is fully orthogonal to mEos3. We then co-expressed a simple amyloid-forming dipeptide repeat, (GA)20 (Chang *et al*., 2016), fused to mEos3.1, to investigate via imaging flow cytometry and DAmFRET its colocalization with Nb condensates and consequent stimulation of amyloid formation. All three nanobody fusions formed puncta as expected, but only mEosNb sequestered the (GA)20-mEos3 fusion (**Figure 1C**) and stimulated its amyloid formation (**Figure 1D**).

### A systematic screening platform

Having demonstrated that mEosNb can direct modifier proteins to mEos3-fused query proteins, we next sought to deploy it systematically across many modifiers and queries. Yeast mating offers a facile and scalable means to do so. Specifically, it allows for the creation of a single parental strain for each modifier and query protein of interest, and then testing them in pairwise combination by arrayed mating (**Figure 2**). We typically express mEos3-fused proteins from *URA3*-marked 2-micron plasmids that have been engineered to maximize copy number variation (Khan *et al*., 2018). To allow for simultaneous selection of mEos3- and mEosNb-fusion plasmids in diploids, we created a *LEU2*-marked 2-micron backbone to express modifiers as joint fusions to mEosNb and BDFP1.6:1.6. Two-micron plasmids are spontaneously lost in 1-5% of daughter cells during cell division, leading to a stable population of 10-30% plasmid-free cells even under selection in liquid media (Christianson *et al*., 1992). By gating our diploid cells into BDFP1.6:1.6-positive and -negative populations, this feature provides an internal negative control for the effect of any modifier (**Figure S1A-B**).

**Figure 2.**
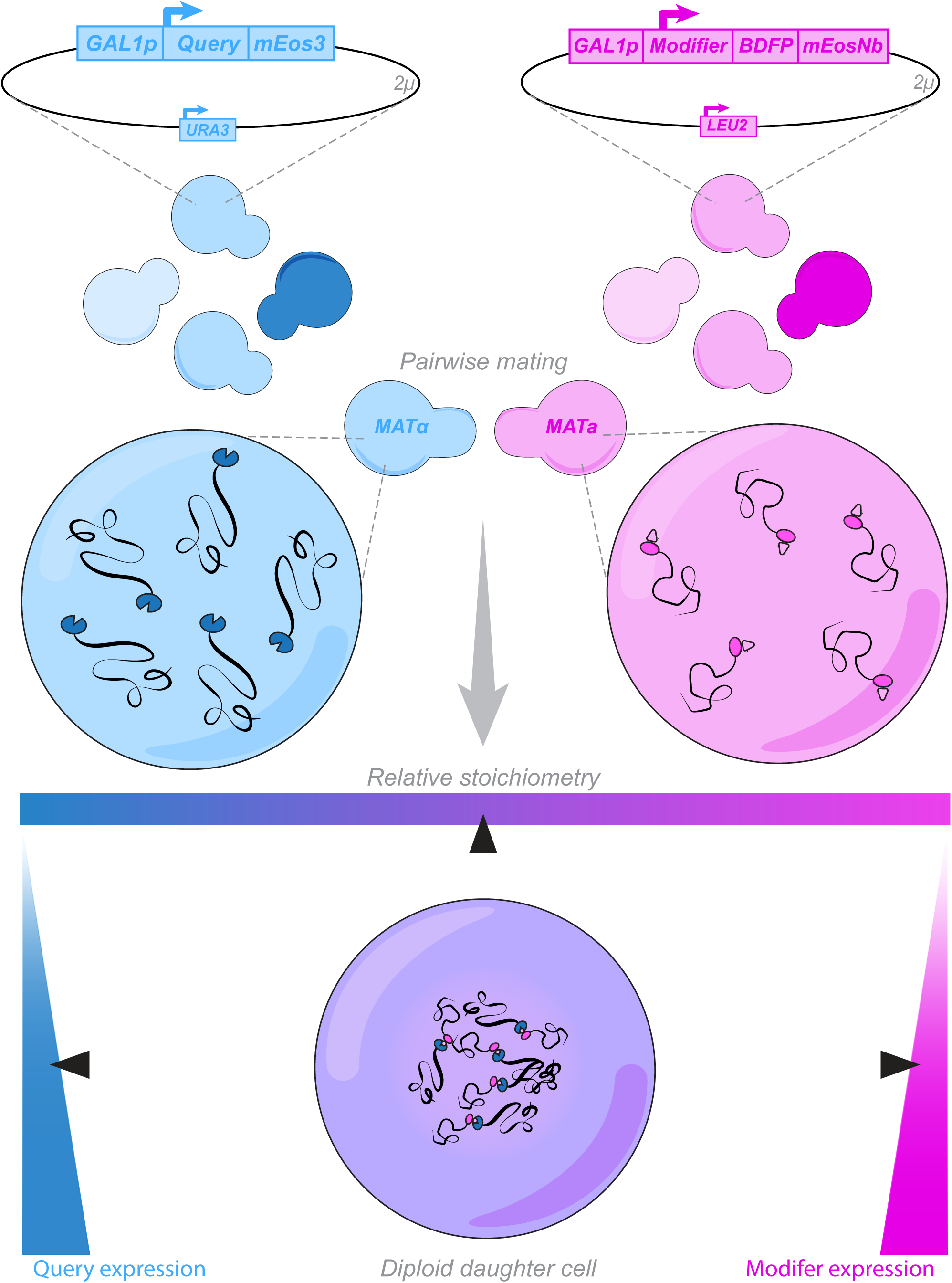
A systematic screening platform. Query and modifier parental strains are developed through plasmid transformation and proteins of interest are inducibly expressed at a dynamic range of intracellular concentrations via a variable copy number 2-micron plasmid. Diploid daughter cells are created through pairwise matings of query and modifier parental strains, creating populations of daughter cells with independently varied expression of query and modifier.

### Application to study coacervation

To demonstrate the utility of our screening platform, we used it to explore the effect of multivalency by modulating the number of mEos3 or mEosNb units from 1 to 4 or 1 to 6, respectively. When expressed on their own, we found that 1×-4× mEos3 remained fully diffuse (**Figure S1C**), consistent with the demonstrated monomericity of this fluorescent protein (Zhang *et al*., 2012; Cranfill *et al*., 2016). We also observed that AmFRET increased with valence from 2× to 4× (**Figure S1D**), but was largely concentration-independent, consistent with the expected energy transfer between intramolecular copies of mEos3. mEosNb proved to be less soluble, forming puncta in a concentration- and valency-dependent manner indicating a weak tendency to self-interact (**Figure S1E**).

We then mated the two series to create all 24 pairwise combinations. We observed via confocal microscopy (**Figure 3A**) and DAmFRET (**Figure 3B**) puncta formation and phase behavior as a function of the expression level and valency of both constructs. The DAmFRET plots revealed three discrete cell populations (see methods for population classification), whose sizes and positions differed with the valence of either construct. Comparing the two datasets provided mechanistic insight by relating the different populations to the presence or absence of puncta. AmFRET increased uniformly with the concentration and valence of both mEos3 and mEosNb (**Figure 3C**, gray populations), even while remaining diffuse well beyond the concentration of mEosNb that assembled when expressed on its own. Hence, mEos3 solubilizes multivalent mEosNb puncta, suggesting that binding to mEos3 competes with binding to another mEosNb. AmFRET levels fell back to the level of intramolecular FRET in cells that contained very high concentrations of multivalent mEos3 but relatively low concentrations of mEosNb (**Figure 3C-D**, blue populations), as expected for mEosNb saturation and subsequent accumulation of unbound diffuse mEos3. Finally, AmFRET levels increased sharply in cells expressing very high levels of both multivalent mEos3 and multivalent mEosNb (**Figure 3C-D**, red population). This separate population of cells also uniquely contained puncta, and therefore results from coacervation of the two proteins. As expected, the concentration of mEos3 at which this population appeared fell with increasing mEos3 valence. Altogether these results echo the mass action dependence of phase separation observed for other multivalent multiprotein systems (Li *et al*., 2012; Banani *et al*., 2016; Nandi *et al*., 2022).

**Figure 3.**
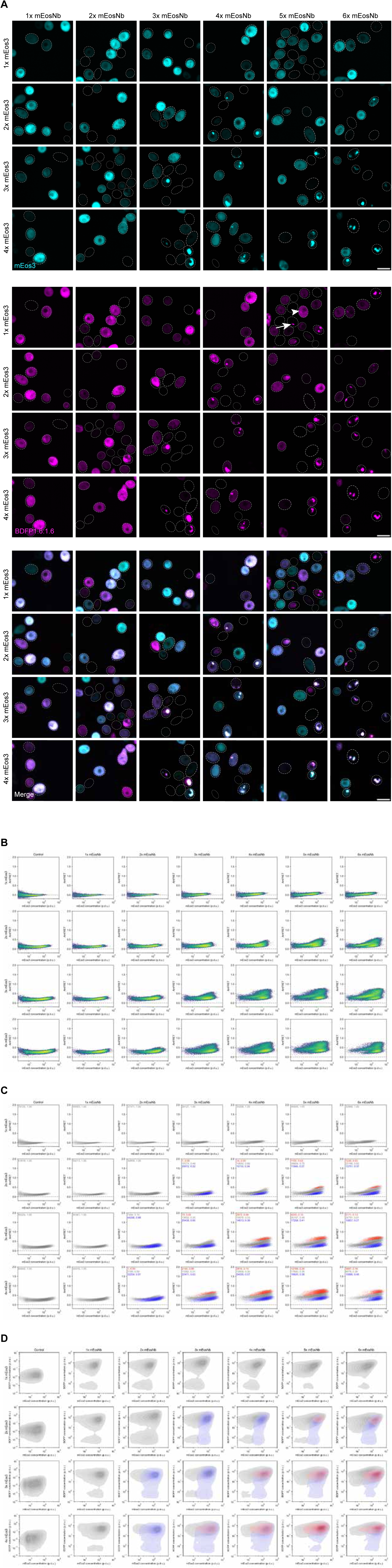
Application to study coacervation. (a) Representative confocal images of diploid cells expressing all pairwise combinations of tandemly repeated mEos3 and mEosNb polypeptides, demonstrating concentration- and valency-dependent puncta formation and colocalization. Punctate multivalent mEosNb (arrow) is solubilized by mEos3 (arrowhead). Image brightness and contrast were individually adjusted to visualize intra-image variation; signal intensity is not comparable between images. Scale bars, 10 μm. (b) DAmFRET plots of pairwise multivalent mEos3 and mEosNb colored by cell density. (c) DAmFRET plots of pairwise multivalent mEos3 and mEosNb colored by population classifications. (d) In vivo phase diagrams constructed from tens of thousands of yeast cells co-expressing mEos3 and mEosNb at varying valencies. Concentration (p.d.u.) was calculated by dividing mEos3 acceptor fluorescence and mEosNb BDFP1.6:1.6 fluorescence by side scatter (SSC), a proxy for cell volume.

### Application to study nucleation mechanisms

We next deployed our multivalent mEosNb series to test the theoretical relationship of nucleation to interactor valence. For this purpose we selected two query proteins that undergo functional nucleation-limited phase transitions and that we have previously analyzed by standard DAmFRET (Khan *et al*., 2018). Both proteins function in innate immune signaling pathways by storing energy in a supersaturated state prior to pathogen-triggered nucleation and polymerization-mediated signal amplification (Saupe, 2020; Gama *et al*., 2024). The structural basis of the functional nucleation barrier appears to be completely different between the two proteins, however. For human ASC, polymerization involves an intermolecular arrangement of stably prefolded subunits into a triple helical polymer (Lu *et al*., 2014). Given the absence of a conformational change between soluble and assembled subunits, ASC nucleation must be determined by a balance of intermolecular interactions and surface energy in accord with classical nucleation theory (Vekilov, 2012; Buell, 2017). Hence, the critical size of the nucleating oligomer will vary with protein concentration, and specifically, will monotonically decrease as concentration increases. In contrast, for SesA from the filamentous fungus *Fusarium haematococcum*, polymerization involves a dramatic conformational change from an intrinsically disordered monomer to a highly ordered amyloid. The many intramolecular degrees of freedom involved with SesA nucleation are not considered in classical nucleation theory, and are expected to reduce the dependence of nucleation on oligomer size (Khan *et al*., 2018). Nevertheless, because each subunit within the amyloid fibril interacts extensively but exclusively with the subunits immediately preceding and succeeding it, we might expect increased nucleation from co-expressed 3× (and higher) mEosNb.

To test these predictions, we mated ASC- or SesA-mEos3 expressing cells with 1× to 6× mEosNb-expressing cells and analyzed the proteins’ behaviors as above. Both proteins formed puncta at higher frequencies with increasing mEosNb valence, and the two fluorescent proteins colocalized in all cases (**Figure 4A-B**). Via DAmFRET we found that monovalent mEosNb had negligible effect on nucleation (**Figure 4C-D**), confirming that mEosNb is itself inert. ASC nucleation proved very responsive to multivalent mEosNb (**Figure 4E**), consistent with the absence of a rate-limiting conformational change. It also increased with valence across the whole series, occurring at lower concentrations as valence increased, confirming the prediction of classical nucleation theory. SesA nucleation proved to be much less responsive to multivalent mEosNb (**Figure 4E**), as predicted. Also consistent with predictions, bivalent mEosNb had no effect, while 3× and higher mEosNb valencies slightly increased nucleation, and all to a comparable extent, consistent with nucleation comprising a rare trimeric folding event.

**Figure 4.**
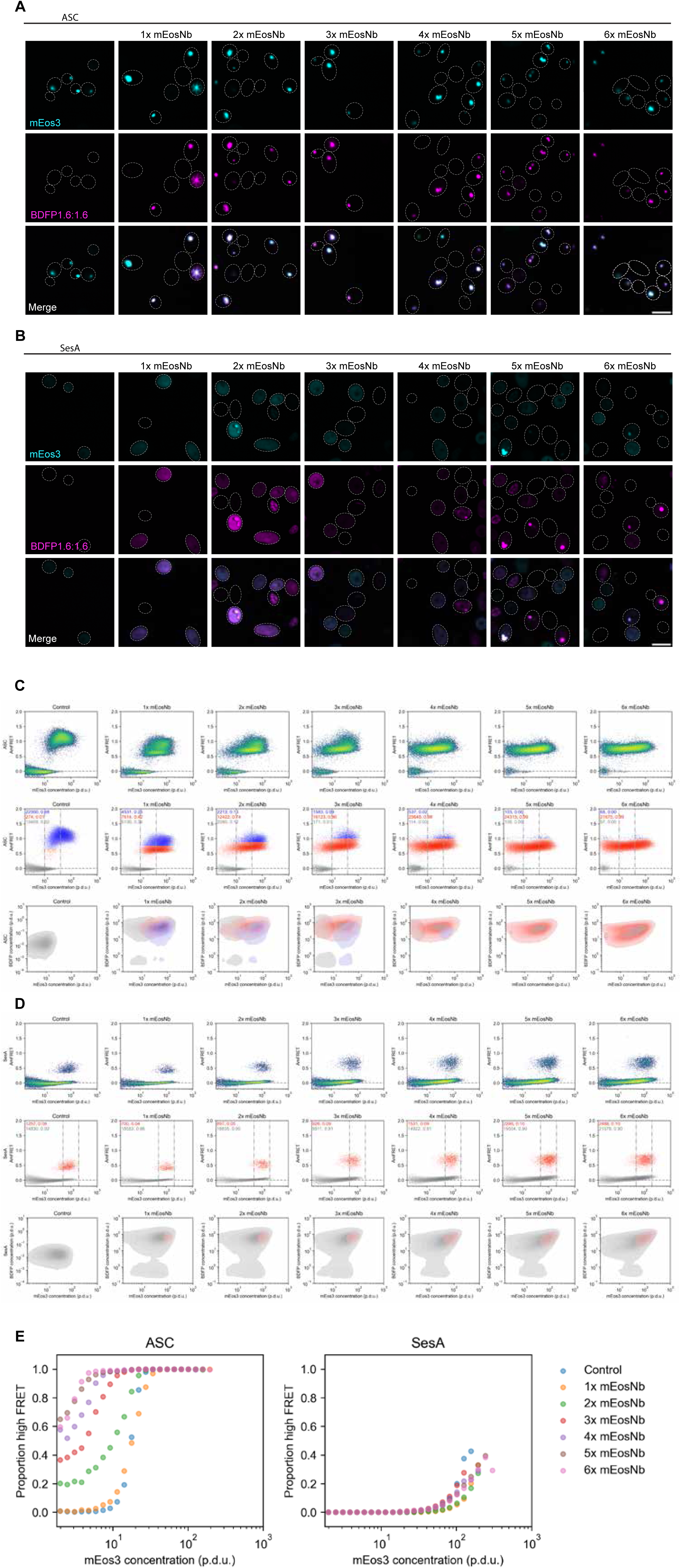
Applications to study nucleation mechanisms. (a) Representative confocal images of query proteins ASC and (b) SesA expressed on their own (leftmost column) or co-expressed with multivalent mEosNb modifiers. Scale bars, 10 μm. (c) DAmFRET plots colored by density (top) and population classification (middle) of ASC co-expressed with control HA tag or tandemly repeated mEosNb reveal the emergence and persistence of a reduced high FRET population (red). mEosNb valency reduces saturating concentration required for self-assembly, as evidenced by the reduction in proportion of monomer-only cells (grey) and increase in proportion of assembly-containing cells (blue and red) within the same window of mEos3 concentration (middle). Grey, red, and blue populations occupy distinct yet overlapping regions on phase diagrams (bottom). (d) DAmFRET plots colored by density (top) and population classification (middle top) of SesA co-expressed with control HA tag or tandemly repeated mEosNb display subtle increase in nucleation frequency in response to mEosNb valency. Increasing mEosNb valency from 2× to 3× markedly increases nucleation, as evidenced by increased proportion of high FRET within the same window of mEos3 concentration, whereas nucleation frequency plateaus at higher mEosNb valencies (middle). Grey and red populations occupy completely overlapping regions on phase diagrams (bottom). Cell counts displayed in population classification plots represent counts and proportions within the concentration window of mEos3. (e) Proportion high FRET cells for ASC (left) and SesA (right) at logarithmically binned intervals of mEos3 concentration demonstrating nucleation mechanism-specific sensitivities to multivalent mEosNb.

## Discussion

We here extended our previously developed method, DAmFRET, for enhanced study of the mechanisms by which proteins modulate phase separation in trans. Our approach exploits the fact that all proteins analyzed by DAmFRET are necessarily fused to the fluorescent protein mEos3, by co-expressing candidate modifier proteins as fusions to a high affinity mEos3-binding protein, mEosNb, that we developed for this purpose. We demonstrated a facile and scalable mating-based approach to test combinations of query and modifier proteins, and deployed it to systematically probe the role of valence in LLPS as well as the nucleation of ordered solid phases.

We demonstrated that mEos3-fused proteins can be recruited to other cellular structures by genetically tagging those structures with mEosNb, and that mEosNb itself does not perturb the behavior of mEos3-fused query proteins. In the case of a simple amyloid forming sequence, recruitment to a mEosNb-fused condensate dramatically accelerated nucleation. Moving forward, mEosNb could be fused to a variety of subcellular targeting sequences. For example, fusions to a nuclear localization or nuclear export signal (NLS or NES, respectively) would facilitate systematic interrogation of phase separation in the nuclear versus cytoplasmic compartments. Fusing mEosNb to other phase separating or aggregating proteins would eliminate colocalization as a variable in studies of heterogenous nucleation, thereby allowing for different contributions of sequence to be teased apart.

We formed entirely synthetic complex coacervates by expressing polypeptides containing up to six mEosNb tandem repeats and up to four mEos3 tandem repeats. The former can be used to scaffold the homo-oligomerization of any mEos3-fused protein of interest, thereby providing a simple way to systematically test the effect of valence on saturating concentrations and kinetics of phase separation in living cells.

Our system is a valuable addition to the existing toolbox of designer condensates (Qian *et al*., 2022; Dai *et al*., 2023). Most such tools repurpose intrinsically disordered regions (IDRs) and other natural multivalent proteins, which have the potential to interact with endogenous cellular factors. Our system avoids such crosstalk by using just two, fully synthetic, well-folded components. With its exceptional solubility and unique ability to report on self-interaction in the soluble phase (via AmFRET), mEos3 is a superior fluorescent reporter of phase behavior. Moreover, it allows for FACS-based high throughput screens of pooled genetic perturbations, which streamline the identification of both sequence and cellular determinants of phase separation. Our development here of mEosNb further increases the functionality of mEos3 by allowing the experimentalist to now specify its subcellular localization or recruit to it any protein activity of interest to interrogate its effect on phase behavior.

## Materials and Methods

### mEos3.1 cloning, expression and purification

A gBlock (IDT) encoding mEos3.1 codon optimized for *E. coli* was cloned into a pET-Avitag-His6 backbone derived from Addgene plasmid 29725 using standard Gibson cloning methods. BL21 (DE3) *E. coli* were transformed with pET-Biotin-His6-mEos3.1 and pET21d-BirA, which encodes the biotin ligase that targets the AviTag and grown overnight in LB containing kanamycin (50 ug/mL) and ampicillin (100 ug/mL). Cells were next diluted to an 0D600 of 0.1 and cultured at 37C until the OD600 was 0.6. The cells were cooled to 25C, and expression of mEos3.1 was induced with 1 mM IPTG for 16 hours at 25C. Cells were collected by centrifugation and resuspended in 100 mL of lysis buffer (20 mM Tris, pH 8.0; 150 mM NaCl, 5 mM beta-mercaptoethanol (BME), 1X cOmplete protease inhibitor cocktail (Roche), 0.25 mg/mL lysozyme). Cells were incubated on ice for 30 minutes, followed by 6 rounds of sonication for 30 seconds each, with 2 minutes on ice between rounds. Lysed cells were centrifuged for 30 minutes at 30,000 x rpm in a 45Ti rotor at 4C. The clarified lysate was applied to 5 mL of Ni-NTA agarose (Qiagen) in a gravity column. The resin was washed with 100 mL of high salt wash buffer (20 mM Tris, pH 8.0; 500 mM NaCl, 20 mM imidazole, 5 mM BME). The resin was then washed with 50 mL of lysis buffer lacking protease inhibitors, and mEos3.1 was eluted with lysis buffer containing 250 mM imidazole. Peak fractions were pooled and dialyzed overnight at 4C against 1X PBS and then supplemented with 10% glycerol and snap-frozen in liquid nitrogen for storage at -70C. The protein was later thawed and diluted into 20 mM Tris 8.0 to lower the ionic strength before application to a 1 mL HiTrap Q HP anion exchange column attached to an AKTA Start FPLC system (Cytiva). Unbound protein was washed away with 20 mM Tris, pH 8.0, and bound proteins were eluted with a linear gradient from 20 mM Tris 8.0 to 20 mM Tris, pH 8.0, containing 1 M NaCl. The peak fractions containing highly pure mEos3.1 were pooled and buffer exchanged into PBS using a 10k MWCO centrifugal concentrator filter (Millipore Amicon Ultra), supplemented with 10% glycerol and snap frozen as described. Biotinylation of mEos3.1 was verified by streptavidin pull-downs. Purification of ALFA-tagged mEos3.1 was carried out as described for untagged mEos3.1, except that the anion exchange step was omitted.

### Yeast strains

Yeast strains were created using standard lithium-acetate transformation as follows. The nanobody display library, clones derived from it, and yeast strain BJ5465, were obtained from (McMahon *et al*., 2018). The gBlock (IDT) encoding NbALFA was cloned into the CEN plasmid derived from the McMahon et al. nanobody library using standard Gibson cloning techniques. We used PCR-based mutagenesis (Goldstein and McCusker, 1999) to first convert the hphMX cassette in rhy1713 (Khan *et al*., 2018) to kanMX using pFA6a-link-yomTagBFP2-Kan (Lee *et al*., 2013) as a template, yielding yeast strain rhy2849. We then created yeast strain rhy2925 by integrating *GAL1*-driven *HMX1* at the endogenous *HIS3* locus using AfeI-linearized rhx3372 (Miller *et al*., 2023). We then created rhy2926 by replacing the *HO* locus with a cassette consisting of: natMX followed by the doxycycline-repressible *tetO7* promoter and counterselectable *URA3* ORFs derived from *C. albicans* and *K. lactis*, followed by a stop codon and coding sequences for µNS 471-721 (Schmitz *et al*., 2009) and BDFP1.6:1.6 (Ding *et al*., 2018). Yeast strains rhy2984 and rhy2985 were made by replacing the *URA3* ORFs with stopless yeast codon-optimized ORFs encoding NbALFA or mEosNb, respectively. Successful integration was confirmed using fluorescence microscopy to observe far red fluorescent puncta when grown in the absence of doxycycline. Yeast strains rhy2054 (*MATα*) and rhy3077 (*MATa*) were created by sequentially mating and sporulating rhy1713 (Khan *et al*., 2018) with the *PDR5* and *ATG8* knockout strains from the *MATa* deletion collection (Open Biosystems). Yeast strains rhy2145 and rhy3279 were made by passaging rhy2054 and rhy3077, respectively, four times on YPD plates containing 3 mM GdHC to eliminate the amyloid form of Rnq1 (Ferreira *et al*., 2001).

### Anti-mEos3 nanobody screening

All steps were derived from methods described in McMahon et al. In the first campaign, a MACS enrichment step (1 µM antigen, anti-biotin microbeads (Miltenyi)) was followed by 3 rounds of FACS (1 µM, 1 µM, and 250 nM antigen). In the second screen, MACS was omitted, and 5X the entire library (500 x 10^6^ cells) was subjected to sorting (1 µM antigen), followed by two additional rounds of FACS (200 nM and 40 nM antigen). Briefly, cells were first grown overnight at 30°C in SD-Trp media. This and all subsequent media contained 100 U PenStrep. The MACS and whole-library FACS steps were initiated by thawing a single library stock containing 2.5 x 10^9^ cells and adding the cells to 1 L SD-Trp media. Subsequent FACS steps had variable cell and culture volumes that depended upon the estimated library diversity, but at least 10X diversity was maintained in samples throughout the screening campaign. For each step, cells were counted on an EC800 cytometer (Sony), and enough cells were pelleted and washed with Sgal-Trp media to resuspend in the desired final volume at a density of 5 x 10^6^ cells/mL and then cultured in Sgal-Trp at 25C for 48 hours to induce nanobody expression. Induced cells were collected by centrifugation and washed twice with Selection Buffer (20 mM HEPES, pH 7.5; 150 mM NaCl, 5 mM Maltose, 0.1% BSA). For MACS enrichment of mEos3.1 binders, cells were first subjected to a depletion step with anti-biotin microbeads and an LD MACS column (Miltenyi) to remove bead-binding nanobodies. Cells were then positively selected by first incubating with mEos3.1 (1 µM), followed by anti-biotin microbeads and an LS MACS column. Enriched cells were collected and cultured in SD-Trp media to expand the cells for glycerol stocks and subsequent screening steps. For FACS, induced cells were resuspended and incubated with mEos3.1 at the desired concentration, along with an anti-HA antibody conjugated to Alexa Fluor 647 (ThermoFisher) at a concentration of 0.5 µM to stain the HA-tagged nanobodies, and incubated with gently mixing. Unbound antibody and mEos3.1 were removed by washes with Selection Buffer, and cells were then resuspended in Selection Buffer containing 1 µg/mL propidium iodide (PI) to stain dead cells. Cells were sorted using a BD Influx cytometer outfitted with a 70-µm tip and operating at 45 psi in 1X PBS. Live, single cells were sorted with a gate derived from the pattern of NbALFA-displaying yeast stained with anti-HA-AF647 and ALFA-tagged mEos3.1.

### mEos3Nb purification

The codon optimized gBlock (IDT) encoding mEosNb was cloned into pET26b with a C-terminal Avitag using standard Gibson cloning techniques. Protein expression was conducted as described above. 2L of cell culture were pelleted and prepared for periplasmic extraction by resuspending them in 100 mL of 0.2 M mM Tris, pH 8.0; 0.5 M sucrose, 0.5 mM EDTA, 1X Roche cOmplete protease inhibitor cocktail. The cells were stirred for 45 minutes at room temperature and then osmotically shocked with 200 mL of ice cold water. The cells were stirred for an additional 45 minutes before adjusting the buffer to 150 mM NaCl, 2 mM MgCl2, and 20 mM imidazole. Cells were centrifuged at 20,000 x g for 20 minutes in a JLA14 rotor at 4°C. The lysate was retrieved and applied to 6 mL of Ni-NTA in a gravity column. The resin was washed with 50 mL of high salt buffer (20 mM HEPES, pH 7.5; 500 mM NaCl, 20 mM imidazole). Next the resin was washed with 50 mL of low salt buffer (20 mM HEPES, pH 7.5; 100 mM NaCl, 20 mM imidazole) and then bound proteins were eluted with the same buffer containing 300 mM imidazole. Peak fractions were collected and dialyzed into low salt buffer, supplemented with glycerol to 10% and snap frozen in liquid nitrogen.

### Biolayer interferometry

mEosNb-3C-Avitag-His protein was incubated with HRV 3C protease (ThermoFisher 88946) overnight at 4°C in a rotator as per manufacturer’s instructions to cleave the 3C-Avitag-His. The reaction was passed through a 5 ml HisTrap column to isolate tag free mEosNb. A biolayer interferometry experiment was performed in the Sartorius Octet R4 instrument in 96-well plates under shaking conditions using the following steps. In Step 1, the Octet® Ni-NTA biosensors (Sartorius 18-5101) were dipped into wells containing PBSi Buffer (1 x PBS supplemented with 15 mM imidazole and 150 mM NaCl) for 5 min for pH equilibration. In Step 2 (Loading step, 10 min), biosensors were moved to wells containing 15 µg/ml Avitag-His-TEV-mEos3.1 for loading followed by step 3 where the biosensors were dipped into PBSi again for 10 min to remove non-specific binding interactions. In Step 4 (Association step, 10 min), the biosensors were dipped into wells containing varying concentrations (15-60 µg/ml) of tag free mEosNb. In step 5 (Dissociation, 10 min), the biosensors were moved to PBSi buffer. The kinetic analysis was performed using Octet Analysis Studio 12.2. The curve fitting analysis was performed assuming 1:1 binding stoichiometry to determine association rate, dissociation rate and dissociation constant (Kd).

### AlphaFold structural predictions

Structural predictions of the mEos3-Nb complex and individual mEos3 and mEosNb multivalent peptides were performed using AlphaFold Server (AlphaFold3) (Abramson *et al*., 2024) using default parameters for the complex and a seed of one for the individual peptides. Predicted structures were visualized and analyzed for structural alignment in ChimeraX-1.6.1 (Meng *et al*., 2023). pLDDT of mEos3 and mEosNb complex is colored using default alphafold palette. Pseudobonds of predicted contacts as determined by PAE at a paired distance less than or equal to 5Å are colored either black (Figure 1B, top) by the default paecontacts palette (Figure 1B, middle).

### DAmFRET preparation and data acquisition

Single parental strains for each modifier and query protein of interest were made via standard lithium-acetate protocol using yeast strains rhy2145 (MATα) and rhy3279 (MATa), respectively. Transformants were grown on standard synthetic media (SD) with either uracil or leucine dropouts for query and modifier, respectively. Single colonies were picked in biological triplicate into Axygen round bottom 96-well microplates containing 200 μL of standard synthetic media containing 2% dextrose. Untransformed rhy2145 and rhy3279 were inoculated in triplicate into complete synthetic media (SD-complete) to serve as color compensation dark controls. Cells were incubated at 30°C, shaking on Heidolph Titramax vibrating platform at 1000 rpm overnight. Cells were spun down and resuspended in synthetic induction media containing 2% galactose with corresponding selection dropout (Sgal-Ura, Sgal-Leu) and 100μM of proteasomal inhibitor peptide MG-132 resuspended in DMSO. Color compensation dark control samples were resuspended in 200 μL Sgal-complete. Cells were returned to 30°C shaking incubation for 16 hr, then were spun down and resuspended in fresh Sgal-Ura or Sgal-Leu and 100μM MG-132 4 hr prior to photoconversion. Microplates containing yeast cells were photoconverted using OmniCure S2000 Elite fitted with a 320–500 nm (violet) filter and a beam collimator (Exfo), positioned 45 cm above the plate, for a duration of 5 min while shaking at 1000 rpm. Immediately following photoconversion, yeast cells were assayed on a nonimaging flow cytometer (BioRad ZE5 Analyzer). 20μl of cells were collected per sample, flow speed 1.8μl/sec and 1.25 sec wash between samples. Samples were periodically shaken during runtime to prevent cell settling. Donor fluorescence was collected from 525/35 emission channels with 488 nm laser excitation; FRET signal was collected from the 593/52 emission channel with 488 nm laser excitation; acceptor fluorescence was collected from the 589/15 emission channel with 561 nm laser excitation; BDFP fluorescence was collected from the 670/30 emission channel with 640 nm excitation; autofluorescence signal was collected from the 460/22 emission channel with 405 nm laser excitation. Imaging and nonimaging flow cytometry of nanobody integrated yeast stains was conducted as described above and in previous work (Khan *et al*., 2018; Venkatesan *et al*., 2019) and image contrast adjustment was performed individually per strain. Manual compensation was performed using non photoconverted mEos3.1 donor and BDFP single color controls, dark control samples, and dsRed2 as a proxy for red mEos3.1. To correct for signal intensity as a function of cell volume, acceptor fluorescence and BDFP fluorescence signals were divided by side scatter (SSC) -- a proxy for cell volume -- to calculate concentration in arbitrary procedure defined units (p.d.u.) (Miller *et al*., 2023).

### DAmFRET data analysis – parental strains

Data analysis and visualization was performed using FSC Express 7 software where biological triplicates were assessed for similarity with regard to DAmFRET profile for query proteins, and BDFP expression for modifier proteins. Within biological triplicate, the highest BDFP expressor (most cells with positive fraction BDFP) was chosen to be the parent for mating, whereas the first triplicate for each query protein was chosen as the parent, since all triplicates had high similarity with regard to acceptor expression and AmFRET.

### Coexpression of query and modifier via mating

Parental query and modifier strains were mated pairwise using 10μl of overnight growth of each parent into 80μl of YPD. Strains were mated at 30°C with shaking on a Heidolph Titramax vibrating platform for 90 min. Cells were resuspended and 10μl of culture was plated onto double selection standard synthetic media (SD-Ura-Leu) agar and was incubated for 2 days at 30°C. Mated cells were inoculated into liquid double selection media containing 2% dextrose, and induction of ectopic protein expression was performed as with single parental strains, while maintaining double selection through data collection.

### DAmFRET data analysis

Live, single yeast cells were gated in FCS Express 7 7.22.0031 (De Novo Software) using forward scatter (FSC), side scatter (SSC), and the 460/22 emission channel (FSC 488/10-A, FSC 488/10-H, SSC 488/10-A, 460/22-405 nm-A) to differentiate these cells from cellular debris and cells with high autofluorescence. Cells were screened for mEos3.1 donor and acceptor fluorescence relative to single color compensation controls and untransformed dark cells. Donor+/Acceptor+ cells were further classified based on BDFP1.6:1.6 signal in Python. Cells with BDFP intensity below 5.6 were considered BDFP1.6:1.6-negative, and cells with BDFP intensity above 10 were considered BDFP1.6:1.6-positive. For wells expressing 1× to 6× mEosNb, only the BDFP1.6:1.6-positive population was used to analyze the effects of mEosNb valency. Population gating for ASC-mEosNB multivalency series was manually performed in FCS Express 7, where gates were drawn based on diploid cells harboring both the ASC-mEos3 plasmid and a second, effectively empty plasmid (encoding an HA tag instead of mEosNb-BDFP1.6:1.6), and adjusted to follow the natural density differences between high FRET populations. Plots in the mEos3-mEosNb valency series had populations assigned based on lines which split the populations in the 6× mEosNb plot for a specific level of mEos3 valency. Plots which did not show evidence of multiple populations had all cells assigned to the same label as the population with rising AmFRET visible at high mEosNb valency. Plots for SesA were divided into two populations, where the high FRET population had both high mEos3 expression and AmFRET. Boundaries for visualizing the change in nucleation for SesA and ASC were selected based on the expression levels with the highest overlap in populations for 1× mEosNb plots. Proportion high FRET plots were generated by calculating the proportion of nucleated cells within 30 logarithmically spaced mEos3 expression bins, only including data from bins with more than 75 cells. Analysis and visualization were performed with Python 3.9.18 and the following packages: fcsparser, matplotlib, seaborn, mpl-scatter-density, pandas, and NumPy. Scripts used to analyze and visualize DAmFRET results are available at https://github.com/jacobnjensen1/Kimbrough-mEosNb-preprint.

### Confocal microscopy

Cells were prepared for imaging using the DAmFRET protocol as described above, with the exception of photoconversion. Post a 4 hr media refresh period, cells are immediately imaged using a high throughput confocal microscope. Images were acquired with an Andor Xyla x4 sCMOS camera at full resolution on a PerkinElmer Opera Phenix spinning disk microscope. Samples were illuminated with 640nm (17mw) and 488nm (20mw) lasers (LUNV 6-line Laser Launch). Emissions filters used to acquire these images were 650-760nm and 500-550nm bandwidth emission filters. A PerkinElmer 63× water objective lens (N.A. 1.15) was used to acquire the images. Z stacks were acquired for a total distance of 0.8μm, with a spacing of 0.2μm. After acquisition, each channel was max projected (NumPy) to combine all images from different planes. Brightness and contrast were adjusted based on the expected brightest biological sample, and those settings were propagated to all other images within the panel, as in Figure 4A-B and Figure S1C, E-F. Due to the large variation in signal intensity, brightness and contrast were individually adjusted on each image to visualize dim signal that otherwise would not be visualized, as in Figure 3A. Cells were identified using the brightfield image and cell outlines were approximately hand drawn using the ellipse tool. Analysis was performed in Fiji version 2.14.0.

## Author Contributions

**Hannah Kimbrough:** investigation; supervision; visualization; writing – original draft; writing – review and editing. **Jacob Jensen**: data curation; formal analysis; software; visualization. **Caleb Weber:** investigation. **Tayla Miller:** investigation. **Cindy Maddera:** investigation. **Vignesh Babu:** investigation. **William B. Redwine:** investigation; supervision; writing – original draft. **Randal Halfmann:** conceptualization; funding acquisition; supervision; writing – original draft; writing – review and editing.

## Acknowledgments

We thank members of the Halfmann lab for constructive input, and members of the Microscopy, Cytometry, Custom Protein Resources, Automation and PCR Technology, and Media Prep cores of the Stowers Institute for assistance with experiments. This work was performed to fulfill, in part, requirements for HK’s thesis research in the Graduate School of the Stowers Institute for Medical Research. This work was supported by the National Institute of General Medical Sciences of the National Institutes of Health (Award Number R01GM130927, to RH) and the Stowers Institute for Medical Research. The funders had no role in study design, data collection and analysis, or manuscript preparation. The content is solely the responsibility of the authors and does not necessarily represent the official views of the funders.

## Supporting Information

Original data underlying this manuscript can be accessed from the Stowers Original Data Repository at https://www.stowers.org/research/publications/LIBPB-2520.

Supplemental material consists of two files: Figure S1 and Table S1.

**Figure S1. Additional data supporting main figures.** (a) The maximum BDFP1.6-A signal for BDFP1.6:1.6-negative (“bdfp-”) cells and the minimum BDFP1.6-A signal for BDFP1.6:1.6-positive (“BDFP+”) cells are shown in blue and red, respectively. Cells where the BDFP1.6-A signal was compensated to 0 or less are not included in these plots, but comprise 79990 cells for the control plot (86% of cells) and 16424 cells for 4× mEos3 with 6× mEosNb (39% of cells). (b) DAmFRET plots show that the bdfp- portion of the well with 4× mEos3 and 6× mEosNb is very similar to the control plot, while the BDFP+ portion shows differences due to expression of 6× mEosNb. (c). Representative confocal microscopy images of parental strains expressing query proteins 1-4× mEos3 displaying diffuse signal at all valencies and concentrations. (d) DAmFRET plot of multivalent mEos3 peptides revealing increased acceptor intensity and AmFRET as a function of valency, resulting from brighter intracellular signal and intramolecular FRET, respectively. (e) Representative confocal microscopy images of parental strains expressing query proteins 1-6× mEosNb, displaying an increase in puncta formation as concentration and valency increases. (f) Representative confocal images of pairwise combinations of multivalent mEos3 and mEosNb daughter cells as in Figure 3A, with brightness and contrast parameters adjusted the same within a fluorescence channel. Scale bars, 10μm.

**Table S1. Plasmids and protein sequences used in this study.**

