## Supplementary figures and images for "A tool to dissect heterotypic determinants of homotypic protein phase behavior"

### Figure S1

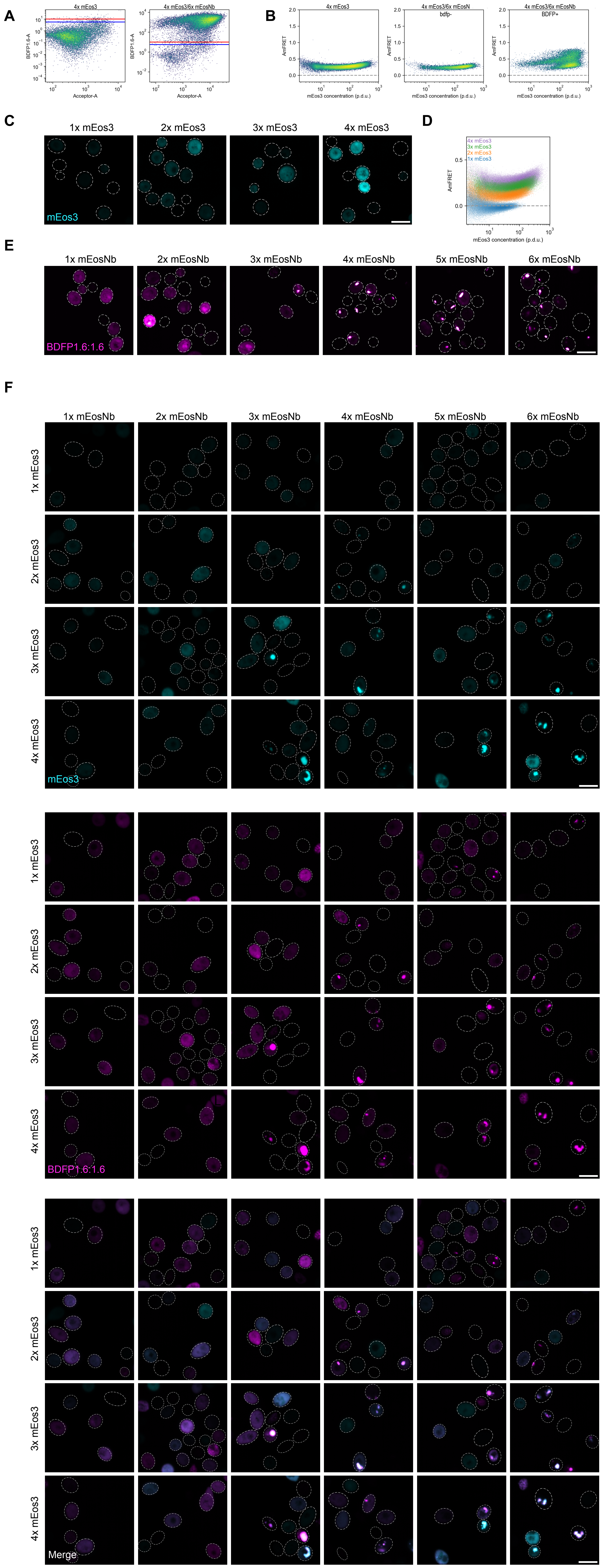
