## Supplementary material for "A tool to dissect heterotypic determinants of homotypic protein phase behavior": Table S1

| Name | Insert | Source | Fusion protein sequence |
| --- | --- | --- | --- |
| rhx3912b | (HA)3 | This study | MTRVSTEQRNLERHMDRILHSPGRHRDSPHSQEHRMGTFHLSLISNSCSPSPLCTRKLGP<br>FCYLLRSLLFCTNL |
| rhx5436 | BDFP1.6:1.6-1xmEosNb | This study | MSMANREVVETLKELLADGEKRVQVAGVIGTNAAEVVKTAVSLLFQEYPELVSPGGCAYTTR<br>RYNMCVRDMNYFLRMCSYAIVAGGASVLDGRMLAGFRDTFNSLGIPLCPAARGIQLMKKI<br>VKEKLATAGMTNIAFVDEPFDYIARVISETEIGHGTGSTGSGSMANREVVETLKELLADGEKR<br>VQVAGVIGTNAAEVVKTAVSLLFQEYPELVSPGGCAYTTRRYNMCVRDMNYFLRMCSYAIV<br>AGGASVLDGRMLAGFRDTFNSLGIPLCPAARGIQLMKKIVKEKLATAGMTNIAFVDEPFDYI<br>ARVISETEIGGGGSMSRDQKMSQVQLQESGGGLVQAGGSLRLSCAASGNISQLVYMGWY<br>RQAPGKERELVAGITHGGITYYADSVKGRFTISRDNAKNTVYLQMNSLKPEDTAVYYCAAFQ<br>WRSDDVGYLNLGPLEYWGQGTQVTVSSKVSGA |
| rhx5437 | BDFP1.6:1.6-2xmEosNb | This study | MSMANREVVETLKELLADGEKRVQVAGVIGTNAAEVVKTAVSLLFQEYPELVSPGGCAYTTR<br>RYNMCVRDMNYFLRMCSYAIVAGGASVLDGRMLAGFRDTFNSLGIPLCPAARGIQLMKKI<br>VKEKLATAGMTNIAFVDEPFDYIARVISETEIGHGTGSTGSGSMANREVVETLKELLADGEKR<br>VQVAGVIGTNAAEVVKTAVSLLFQEYPELVSPGGCAYTTRRYNMCVRDMNYFLRMCSYAIV<br>AGGASVLDGRMLAGFRDTFNSLGIPLCPAARGIQLMKKIVKEKLATAGMTNIAFVDEPFDYI<br>ARVISETEIGGGGSMSRDQKMSQVQLQESGGGLVQAGGSLRLSCAASGNISQLVYMGWY<br>RQAPGKERELVAGITHGGITYYADSVKGRFTISRDNAKNTVYLQMNSLKPEDTAVYYCAAFQ<br>WRSDDVGYLNLGPLEYWGQGTQVTVSSKVSGAGGGGSMSRDQKMSQVQLQESGGGLV<br>QAGGSLRLSCAASGNISQLVYMGWYRQAPGKERELVAGITHGGITYYADSVKGRFTISRDN<br>AKNTVYLQMNSLKPEDTAVYYCAAFQWRSDDVGYLNLGPLEYWGQGTQVTVSSKVSGA |
| rhx5438 | BDFP1.6:1.6-3xmEosNb | This study | MSMANREVVETLKELLADGEKRVQVAGVIGTNAAEVVKTAVSLLFQEYPELVSPGGCAYTTR<br>RYNMCVRDMNYFLRMCSYAIVAGGASVLDGRMLAGFRDTFNSLGIPLCPAARGIQLMKKI<br>VKEKLATAGMTNIAFVDEPFDYIARVISETEIGHGTGSTGSGSMANREVVETLKELLADGEKR<br>VQVAGVIGTNAAEVVKTAVSLLFQEYPELVSPGGCAYTTRRYNMCVRDMNYFLRMCSYAIV<br>AGGASVLDGRMLAGFRDTFNSLGIPLCPAARGIQLMKKIVKEKLATAGMTNIAFVDEPFDYI<br>ARVISETEIGGGGSMSRDQKMSQVQLQESGGGLVQAGGSLRLSCAASGNISQLVYMGWY<br>RQAPGKERELVAGITHGGITYYADSVKGRFTISRDNAKNTVYLQMNSLKPEDTAVYYCAAFQ<br>WRSDDVGYLNLGPLEYWGQGTQVTVSSKVSGAGGGGSMSRDQKMSQVQLQESGGGLV<br>QAGGSLRLSCAASGNISQLVYMGWYRQAPGKERELVAGITHGGITYYADSVKGRFTISRDN<br>AKNTVYLQMNSLKPEDTAVYYCAAFQWRSDDVGYLNLGPLEYWGQGTQVTVSSKVSGAG<br>GGGSMSRDQKMSQVQLQESGGGLVQAGGSLRLSCAASGNISQLVYMGWYRQAPGKER<br>ELVAGITHGGITYYADSVKGRFTISRDNAKNTVYLQMNSLKPEDTAVYYCAAFQWRSDDVG<br>YLNLGPLEYWGQGTQVTVSSKVSGA |
| rhx5439 | BDFP1.6:1.6-4xmEosNb | This study | MSMANREVVETLKELLADGEKRVQVAGVIGTNAAEVVKTAVSLLFQEYPELVSPGGCAYTTR<br>RYNMCVRDMNYFLRMCSYAIVAGGASVLDGRMLAGFRDTFNSLGIPLCPAARGIQLMKKI<br>VKEKLATAGMTNIAFVDEPFDYIARVISETEIGHGTGSTGSGSMANREVVETLKELLADGEKR<br>VQVAGVIGTNAAEVVKTAVSLLFQEYPELVSPGGCAYTTRRYNMCVRDMNYFLRMCSYAIV<br>AGGASVLDGRMLAGFRDTFNSLGIPLCPAARGIQLMKKIVKEKLATAGMTNIAFVDEPFDYI<br>ARVISETEIGGGGSMSRDQKMSQVQLQESGGGLVQAGGSLRLSCAASGNISQLVYMGWY<br>RQAPGKERELVAGITHGGITYYADSVKGRFTISRDNAKNTVYLQMNSLKPEDTAVYYCAAFQ<br>WRSDDVGYLNLGPLEYWGQGTQVTVSSKVSGAGGGGSMSRDQKMSQVQLQESGGGLV<br>QAGGSLRLSCAASGNISQLVYMGWYRQAPGKERELVAGITHGGITYYADSVKGRFTISRDN<br>AKNTVYLQMNSLKPEDTAVYYCAAFQWRSDDVGYLNLGPLEYWGQGTQVTVSSKVSGAG<br>GGGSMSRDQKMSQVQLQESGGGLVQAGGSLRLSCAASGNISQLVYMGWYRQAPGKER<br>ELVAGITHGGITYYADSVKGRFTISRDNAKNTVYLQMNSLKPEDTAVYYCAAFQWRSDDVG<br>YLNLGPLEYWGQGTQVTVSSKVSGAGGGGSMSRDQKMSQVQLQESGGGLVQAGGSLRL<br>SCAASGNISQLVYMGWYRQAPGKERELVAGITHGGITYYADSVKGRFTISRDNAKNTVYLQ<br>MNSLKPEDTAVYYCAAFQWRSDDVGYLNLGPLEYWGQGTQVTVSSKVSGA |
| rhx5440 | BDFP1.6:1.6-5xmEosNb | This study | MSMANREVVETLKELLADGEKRVQVAGVIGTNAAEVVKTAVSLLFQEYPELVSPGGCAYTTR<br>RYNMCVRDMNYFLRMCSYAIVAGGASVLDGRMLAGFRDTFNSLGIPLCPAARGIQLMKKI<br>VKEKLATAGMTNIAFVDEPFDYIARVISETEIGHGTGSTGSGSMANREVVETLKELLADGEKR<br>VQVAGVIGTNAAEVVKTAVSLLFQEYPELVSPGGCAYTTRRYNMCVRDMNYFLRMCSYAIV<br>AGGASVLDGRMLAGFRDTFNSLGIPLCPAARGIQLMKKIVKEKLATAGMTNIAFVDEPFDYI<br>ARVISETEIGGGGSMSRDQKMSQVQLQESGGGLVQAGGSLRLSCAASGNISQLVYMGWY<br>RQAPGKERELVAGITHGGITYYADSVKGRFTISRDNAKNTVYLQMNSLKPEDTAVYYCAAFQ<br>WRSDDVGYLNLGPLEYWGQGTQVTVSSKVSGAGGGGSMSRDQKMSQVQLQESGGGLV<br>QAGGSLRLSCAASGNISQLVYMGWYRQAPGKERELVAGITHGGITYYADSVKGRFTISRDN<br>AKNTVYLQMNSLKPEDTAVYYCAAFQWRSDDVGYLNLGPLEYWGQGTQVTVSSKVSGAG<br>GGGSMSRDQKMSQVQLQESGGGLVQAGGSLRLSCAASGNISQLVYMGWYRQAPGKER<br>ELVAGITHGGITYYADSVKGRFTISRDNAKNTVYLQMNSLKPEDTAVYYCAAFQWRSDDVG<br>YLNLGPLEYWGQGTQVTVSSKVSGAGGGGSMSRDQKMSQVQLQESGGGLVQAGGSLRL<br>SCAASGNISQLVYMGWYRQAPGKERELVAGITHGGITYYADSVKGRFTISRDNAKNTVYLQ<br>MNSLKPEDTAVYYCAAFQWRSDDVGYLNLGPLEYWGQGTQVTVSSKVSGAGGGGSMSR<br>DQKMSQVQLQESGGGLVQAGGSLRLSCAASGNISQLVYMGWYRQAPGKERELVAGITHG<br>GITYYADSVKGRFTISRDNAKNTVYLQMNSLKPEDTAVYYCAAFQWRSDDVGYLNLGPLEY<br>WGQGTQVTVSSKVSGA |
| rhx5441 | BDFP1.6:1.6-6xmEosNb | This study | MSMANREVVETLKELLADGEKRVQVAGVIGTNAAEVVKTAVSLLFQEYPELVSPGGCAYTTR<br>RYNMCVRDMNYFLRMCSYAIVAGGASVLDGRMLAGFRDTFNSLGIPLCPAARGIQLMKKI<br>VKEKLATAGMTNIAFVDEPFDYIARVISETEIGHGTGSTGSGSMANREVVETLKELLADGEKR<br>VQVAGVIGTNAAEVVKTAVSLLFQEYPELVSPGGCAYTTRRYNMCVRDMNYFLRMCSYAIV<br>AGGASVLDGRMLAGFRDTFNSLGIPLCPAARGIQLMKKIVKEKLATAGMTNIAFVDEPFDYI<br>ARVISETEIGGGGSMSRDQKMSQVQLQESGGGLVQAGGSLRLSCAASGNISQLVYMGWY<br>RQAPGKERELVAGITHGGITYYADSVKGRFTISRDNAKNTVYLQMNSLKPEDTAVYYCAAFQ<br>WRSDDVGYLNLGPLEYWGQGTQVTVSSKVSGAGGGGSMSRDQKMSQVQLQESGGGLV<br>QAGGSLRLSCAASGNISQLVYMGWYRQAPGKERELVAGITHGGITYYADSVKGRFTISRDN<br>AKNTVYLQMNSLKPEDTAVYYCAAFQWRSDDVGYLNLGPLEYWGQGTQVTVSSKVSGAG<br>GGGSMSRDQKMSQVQLQESGGGLVQAGGSLRLSCAASGNISQLVYMGWYRQAPGKER<br>ELVAGITHGGITYYADSVKGRFTISRDNAKNTVYLQMNSLKPEDTAVYYCAAFQWRSDDVG<br>YLNLGPLEYWGQGTQVTVSSKVSGAGGGGSMSRDQKMSQVQLQESGGGLVQAGGSLRL<br>SCAASGNISQLVYMGWYRQAPGKERELVAGITHGGITYYADSVKGRFTISRDNAKNTVYLQ<br>MNSLKPEDTAVYYCAAFQWRSDDVGYLNLGPLEYWGQGTQVTVSSKVSGAGGGGSMSR<br>DQKMSQVQLQESGGGLVQAGGSLRLSCAASGNISQLVYMGWYRQAPGKERELVAGITHG<br>GITYYADSVKGRFTISRDNAKNTVYLQMNSLKPEDTAVYYCAAFQWRSDDVGYLNLGPLEY<br>WGQGTQVTVSSKVSGAGGGGSMSRDQKMSQVQLQESGGGLVQAGGSLRLSCAASGNI<br>SQLVYMGWYRQAPGKERELVAGITHGGITYYADSVKGRFTISRDNAKNTVYLQMNSLKPED<br>TAVYYCAAFQWRSDDVGYLNLGPLEYWGQGTQVTVSSKVSGA |
| rhx5092 | mEos3.1 | This study | MSAIKPDMKIKLRMEGNVNGHHFVIDGDGTGKPFEGKQSMDLEVKEGGPLPFAFDILTTAF<br>HYGNRVFAKYPDNIQDYFKQSFPKGYSWERSLTFEDGGICNARNDITMEGDTFYNKVRFY<br>GTNFPANGPVMQKKTLKWEPSTEKMYVRDGVLTGDVEMALLLEGNAHYRCDFRTTYKAKE<br>KGVKLPGAHFVDHCIEILSHDKDYNKVKLYEHAVAHSGLPDNARR |
| rhx3270 | 2xmEos3.1 | This study | MSAIKPDMKIKLRMEGNVNGHHFVIDGDGTGKPFEGKQSMDLEVKEGGPLPFAFDILTTAF<br>HYGNRVFAKYPDNIQDYFKQSFPKGYSWERSLTFEDGGICNARNDITMEGDTFYNKVRFY<br>GTNFPANGPVMQKKTLKWEPSTEKMYVRDGVLTGDVEMALLLEGNAHYRCDFRTTYKAKE<br>KGVKLPGAHFVDHCIEILSHDKDYNKVKLYEHAVAHSGLPDNARRGGGGSSSAIKPDMKIK<br>LRMEGNVNGHHFVIDGDGTGKPFEGKQSMDLEVKEGGPLPFAFDILTTAFHYGNRVFAKY<br>PDNIQDYFKQSFPKGYSWERSLTFEDGGICNARNDITMEGDTFYNKVRFYGTNFPANGPV<br>MQKKTLKWEPSTEKMYVRDGVLTGDVEMALLLEGNAHYRCDFRTTYKAKEKGVKLPGAHF<br>VDHCIEILSHDKDYNKVKLYEHAVAHSGLPDNARR |
| rhx3269 | 3xmEos3.1 | This study | MSAIKPDMKIKLRMEGNVNGHHFVIDGDGTGKPFEGKQSMDLEVKEGGPLPFAFDILTTAF<br>HYGNRVFAKYPDNIQDYFKQSFPKGYSWERSLTFEDGGICNARNDITMEGDTFYNKVRFY<br>GTNFPANGPVMQKKTLKWEPSTEKMYVRDGVLTGDVEMALLLEGNAHYRCDFRTTYKAKE<br>KGVKLPGAHFVDHCIEILSHDKDYNKVKLYEHAVAHSGLPDNARRGGGGSSSAIKPDMKIK<br>LRMEGNVNGHHFVIDGDGTGKPFEGKQSMDLEVKEGGPLPFAFDILTTAFHYGNRVFAKY<br>PDNIQDYFKQSFPKGYSWERSLTFEDGGICNARNDITMEGDTFYNKVRFYGTNFPANGPV<br>MQKKTLKWEPSTEKMYVRDGVLTGDVEMALLLEGNAHYRCDFRTTYKAKEKGVKLPGAHF<br>VDHCIEILSHDKDYNKVKLYEHAVAHSGLPDNARRGGGGSSSAIKPDMKIKLRMEGNVNG<br>HHFVIDGDGTGKPFEGKQSMDLEVKEGGPLPFAFDILTTAFHYGNRVFAKYPDNIQDYFKQ<br>SFPKGYSWERSLTFEDGGICNARNDITMEGDTFYNKVRFYGTNFPANGPVMQKKTLKWEP<br>STEKMYVRDGVLTGDVEMALLLEGNAHYRCDFRTTYKAKEKGVKLPGAHFVDHCIEILSHD<br>KDYNKVKLYEHAVAHSGLPDNARR |
| rhx2912 | 4xmEos3.1 | This study | MSAIKPDMKIKLRMEGNVNGHHFVIDGDGTGKPFEGKQSMDLEVKEGGPLPFAFDILTTAF<br>HYGNRVFAKYPDNIQDYFKQSFPKGYSWERSLTFEDGGICNARNDITMEGDTFYNKVRFY<br>GTNFPANGPVMQKKTLKWEPSTEKMYVRDGVLTGDVEMALLLEGNAHYRCDFRTTYKAKE<br>KGVKLPGAHFVDHCIEILSHDKDYNKVKLYEHAVAHSGLPDNARRGGGGSSSAIKPDMKIK<br>LRMEGNVNGHHFVIDGDGTGKPFEGKQSMDLEVKEGGPLPFAFDILTTAFHYGNRVFAKY<br>PDNIQDYFKQSFPKGYSWERSLTFEDGGICNARNDITMEGDTFYNKVRFYGTNFPANGPV<br>MQKKTLKWEPSTEKMYVRDGVLTGDVEMALLLEGNAHYRCDFRTTYKAKEKGVKLPGAHF<br>VDHCIEILSHDKDYNKVKLYEHAVAHSGLPDNARRGGGGSSSAIKPDMKIKLRMEGNVNG<br>HHFVIDGDGTGKPFEGKQSMDLEVKEGGPLPFAFDILTTAFHYGNRVFAKYPDNIQDYFKQ<br>SFPKGYSWERSLTFEDGGICNARNDITMEGDTFYNKVRFYGTNFPANGPVMQKKTLKWEP<br>STEKMYVRDGVLTGDVEMALLLEGNAHYRCDFRTTYKAKEKGVKLPGAHFVDHCIEILSHD<br>KDYNKVKLYEHAVAHSGLPDNARRGGGGSSSAIKPDMKIKLRMEGNVNGHHFVIDGDGT<br>GKPFEGKQSMDLEVKEGGPLPFAFDILTTAFHYGNRVFAKYPDNIQDYFKQSFPKGYSWER<br>SLTFEDGGICNARNDITMEGDTFYNKVRFYGTNFPANGPVMQKKTLKWEPSTEKMYVRDG<br>VLTGDVEMALLLEGNAHYRCDFRTTYKAKEKGVKLPGAHFVDHCIEILSHDKDYNKVKLYEH<br>AVAHSGLPDNARR |
| rhx0927 | ASC-mEos3.1 | Khan et al., 2018 | MSGRARDAILDALENLTAEELKKFKLKLLSVPLREGYGRIPRGALLSMDALDLTDKLVSFYLE<br>TYGAELTANVLRDMGLQEMAGQLQAATHQGSGAAPAGIQAPPQSAAKPGLHFIDQHRAAL<br>IARVTNVEWLLDALYGKVLTDEQYQAVRAEPTNPSKMRKLFSFTPAWNWTCKDLLLQALRE<br>SQSYLVEDLERSGAEAAAREAAAREAAAREAAARINMSAIKPDMKIKLRMEGNVNGHHFVI<br>DGDGTGKPFEGKQSMDLEVKEGGPLPFAFDILTTAFHYGNRVFAKYPDNIQDYFKQSFPKG<br>YSWERSLTFEDGGICNARNDITMEGDTFYNKVRFYGTNFPANGPVMQKKTLKWEPSTEKM<br>YVRDGVLTGDVEMALLLEGNAHYRCDFRTTYKAKEKGVKLPGAHFVDHCIEILSHDKDYNK<br>VKLYEHAVAHSGLPDNARR |
| rhx1111 | SesA-mEos3.1 | Khan et al., 2018 | MSEATVSELISSTIKSLEEGIRCHKDVEDDNGLREAFHEAGRGLSVVNQALQAAQGNSARE<br>PQSAMNPLRVCNTKANISASIFKAVAQAPKTSRFEGYKEAVRQEGNGQTVEVLVVGMMKD<br>VCALAENFAIQDQVKKLHETIEKLSAMKPSLPKEGPGHTFTNIGNGNQYNATDGPQNINQG<br>GNQVTGGTFPGTVNFGSPWPQNPPFEHRNGAEAAAREAAAREAAAREAAARINMSAIKPD<br>MKIKLRMEGNVNGHHFVIDGDGTGKPFEGKQSMDLEVKEGGPLPFAFDILTTAFHYGNRV<br>FAKYPDNIQDYFKQSFPKGYSWERSLTFEDGGICNARNDITMEGDTFYNKVRFYGTNFPAN<br>GPVMQKKTLKWEPSTEKMYVRDGVLTGDVEMALLLEGNAHYRCDFRTTYKAKEKGVKLPG<br>AHFVDHCIEILSHDKDYNKVKLYEHAVAHSGLPDNARR |
